## Supplementary_Information for "A high-affinity split-HaloTag for live-cell protein labeling"

### Supplementary Figures

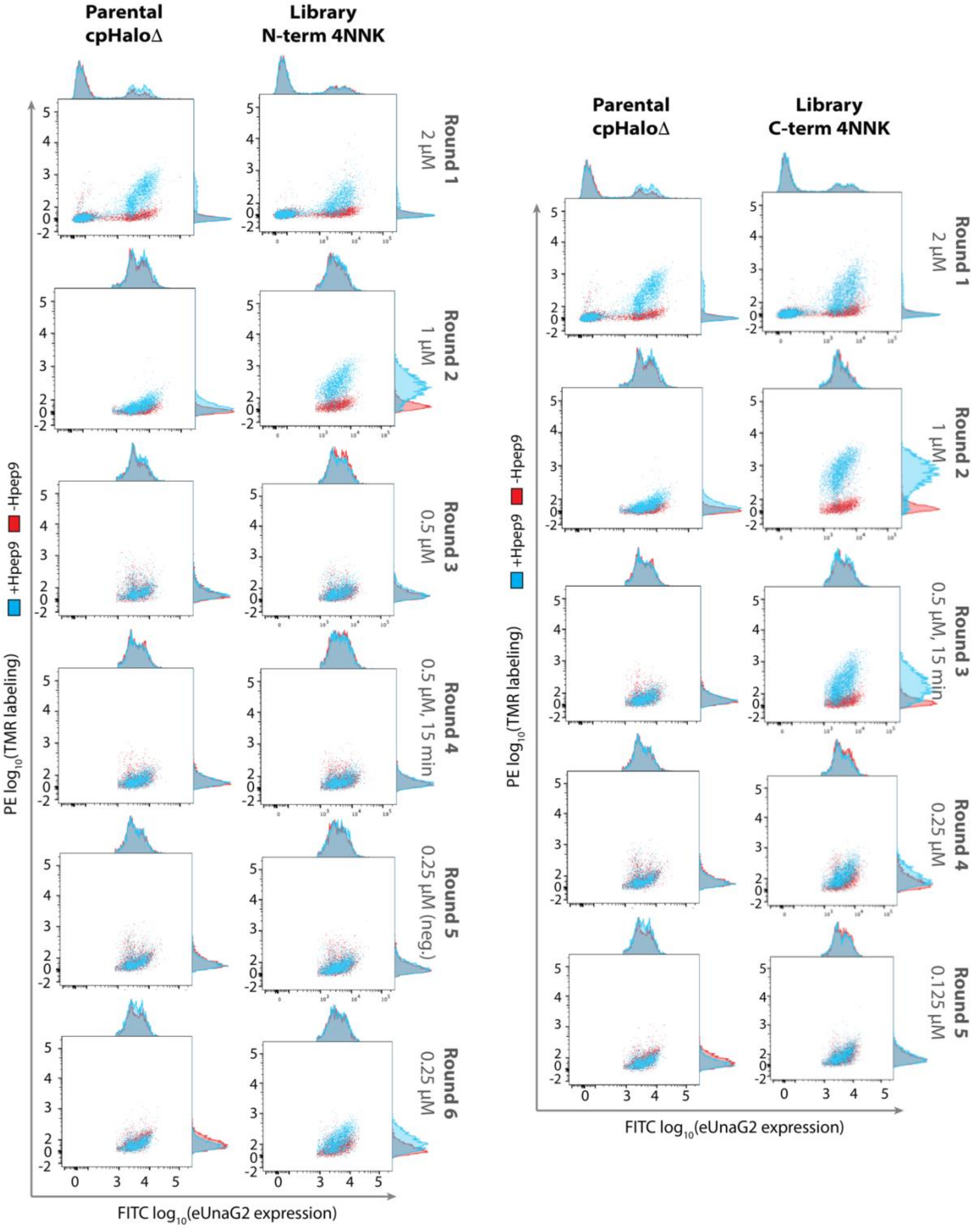

**Supplementary Fig 1.** Selection of cpHaloΔ yeast libraries by FACS.

FACS plots of yeast libraries at each round of selection for cpHaloΔ yeast library with N and C-terminal 4NNK extension. The screening was conducted for 5-6 rounds until no TMR labeling could be detected at the low Hpep9 concentrations.

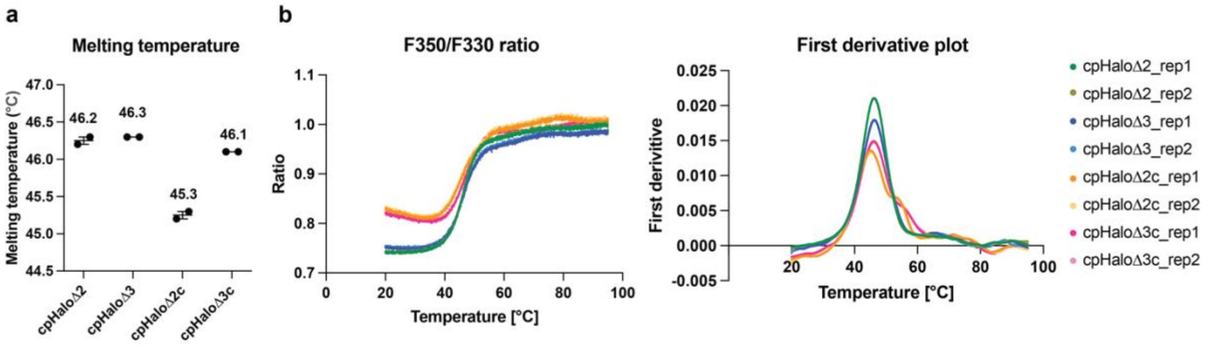

**Supplementary Fig 2.** Melting temperature of cpHaloΔ2 and cpHaloΔ3.

Melting temperature of cpHaloΔ2 and cpHaloΔ3 were determined by nanoDSF. cpHaloΔX stands for His-tagged proteins and cpHaloΔXc represents proteins without His-tag, which were prepared via TEV-cleavage, followed by reverse-IMAC and SEC purification. (**a**), Average melting temperature for each variant from technical duplicate measurements are indicated above each variant. (**b**), Intrinsic fluorescence intensity ratios at 350 nm and 330 nm were plotted as a function of temperature, ranging from 20°C to 95°C (left). First derivative of the ratio was plotted against temperature (right).

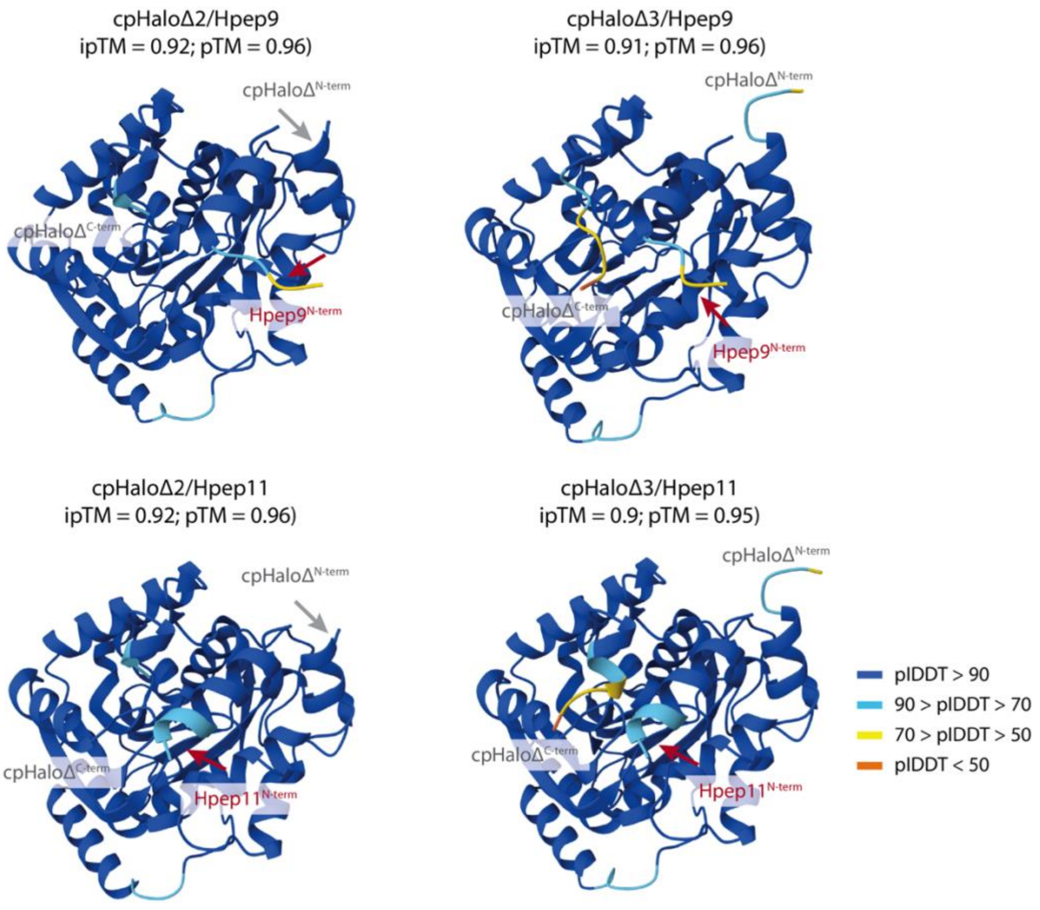

**Supplementary Fig 3.** Split-HaloTag structures predicted by AlphaFold3.

cpHaloΔ/Hpep complex structures were predicted using AlphaFold-Multimer with default parameters. Five models were generated per complex. The interface predicted template modeling (ipTM) and the predicted template modeling (pTM) scores of the top ranked model for each split-HaloTag pair were listed. For visualization, the predicted structures were colored according to the per-atom plDDT values, reflecting the model's confidence in the local structure. Due to the low structural confidence of Hpep9 in complex with both cpHaloΔ2 and cpHaloΔ3, particularly in the N-terminal region of the predicted α-helix, we conducted MD simulations only for the Hpep11 complex.

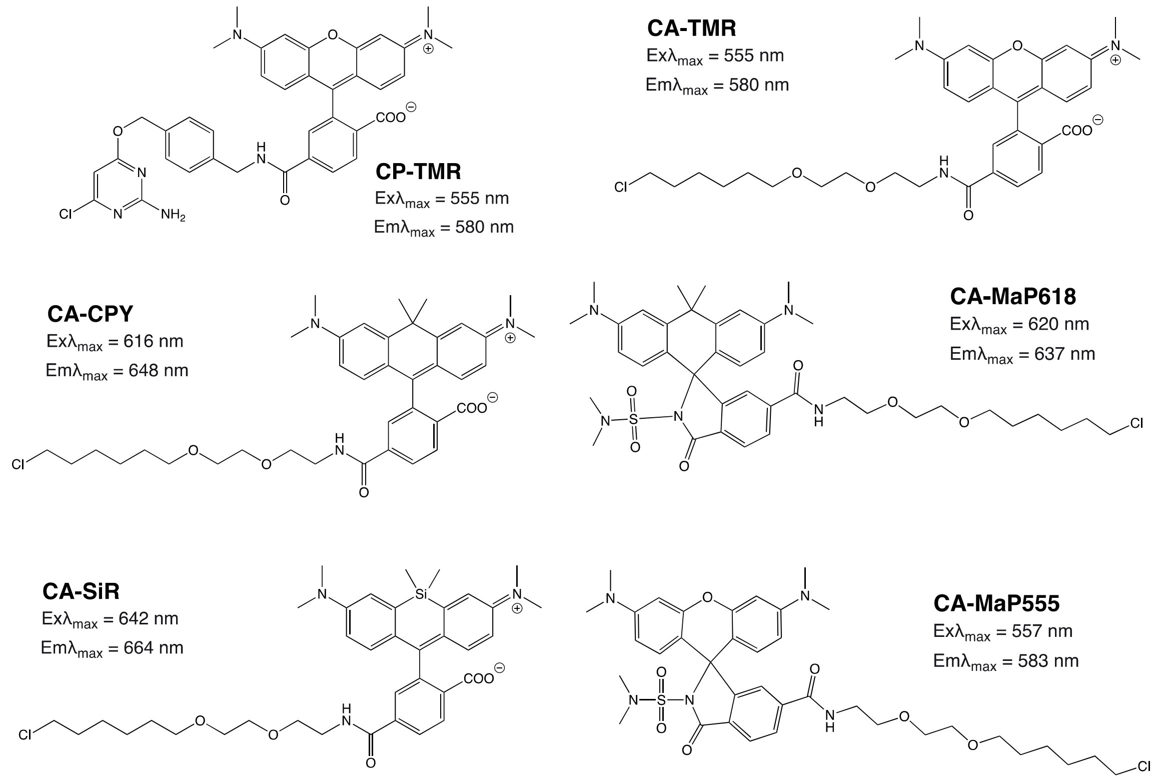

**Supplementary Fig 4.** Fluorescent SLP substrates used in this study.

Fluorescent SNAP-tag (CP-), HaloTag (CA-) substrates and their corresponding wavelengths of excitation/emission maximum.

**a**

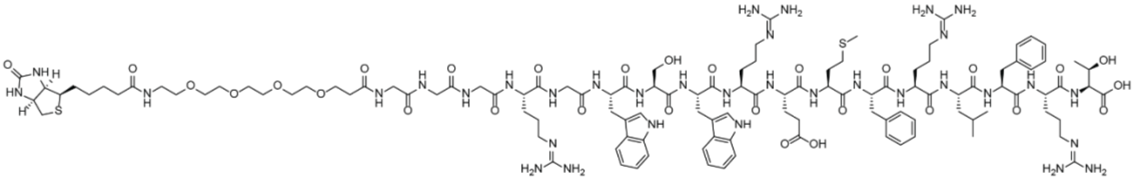

**b**

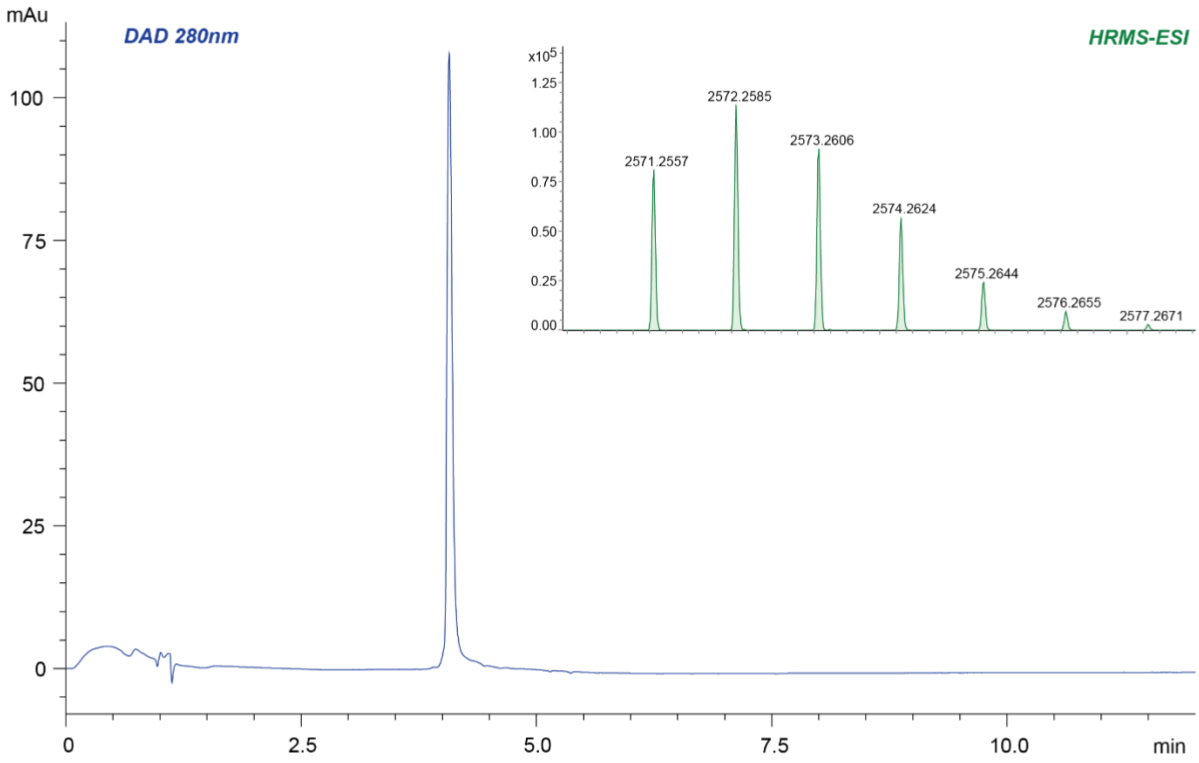

**Supplementary Fig 5.** HPLC-HRMS analysis of the biotinylated Hpep9.

(**a**), Chemical structures of biotinylated Hpep9. (**b**), HPLC-MS analysis of the purified peptide, showing the 280 nm DAD chromatogram and the isotopically resolved intact mass of the corresponding DAD peak. High-resolution mass spectrometry (HRMS, ESI) analysis: calculated mass for $C_{116}H_{174}N_{34}O_{29}S_2$, 2571.2627; observed mass, 2571.2557.

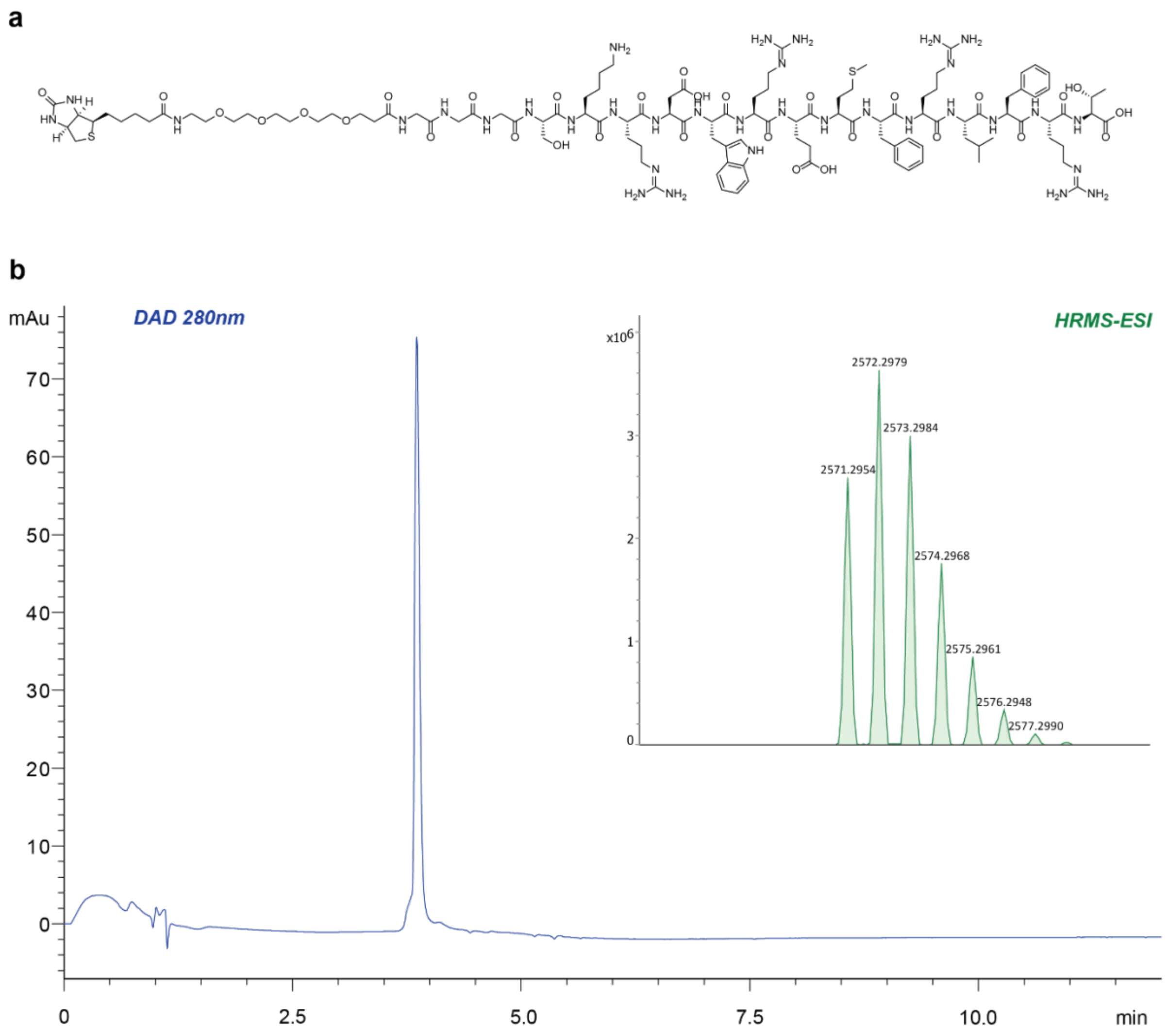

**Supplementary Fig 6.** HPLC-HRMS analysis of the biotinylated Hpep11.

(**a**), Chemical structures of biotinylated Hpep11. (**b**), HPLC-MS analysis of the purified peptide, showing the 280 nm DAD chromatogram and the isotopically resolved intact mass of the corresponding DAD peak. HRMS (ESI) analysis: calculated mass for $C_{113}H_{178}N_{34}O_{31}S_2$, 2571.2839; observed mass, 2571.2954.

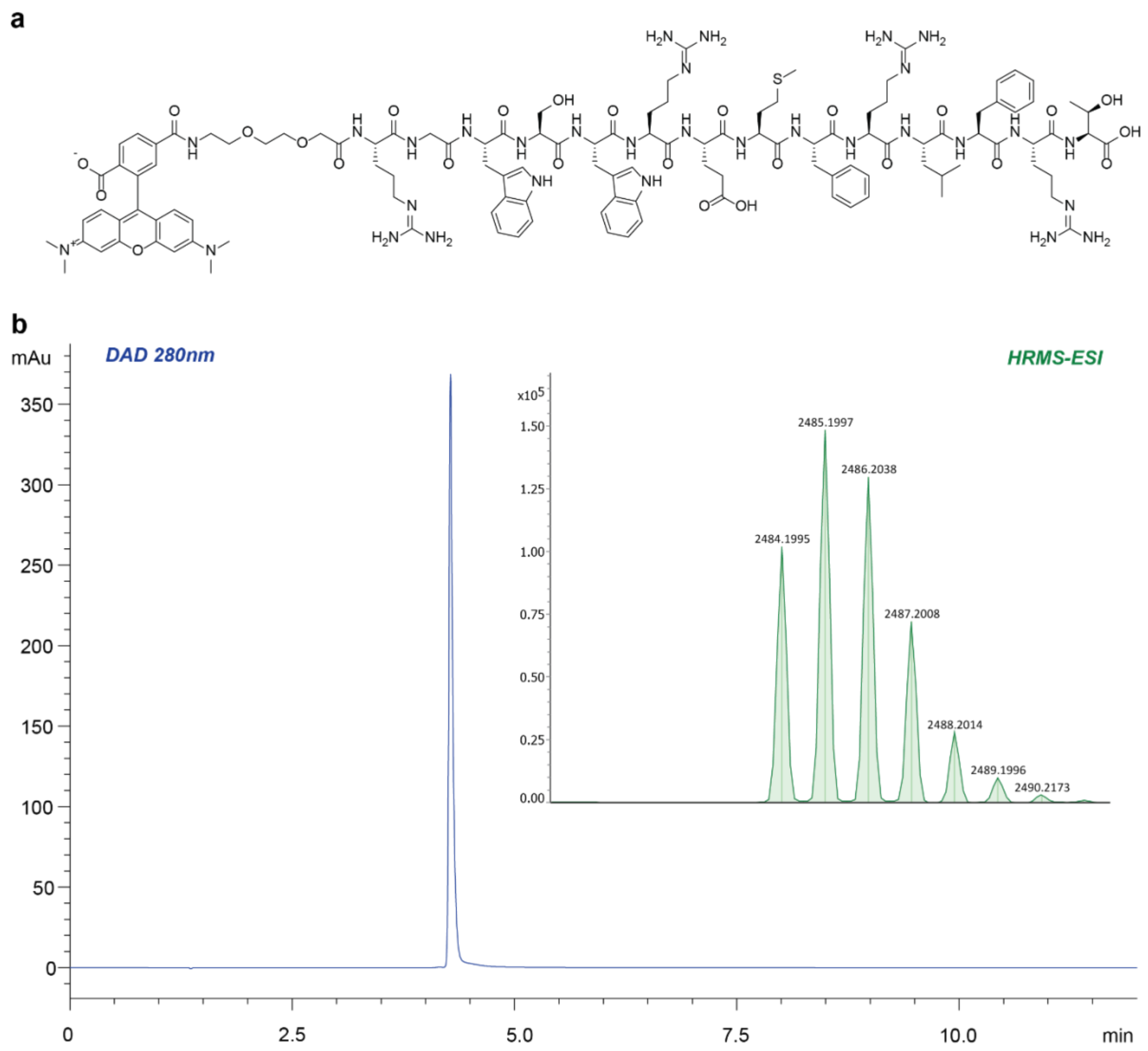

**Supplementary Fig 7.** HPLC-HRMS analysis of the TMR-conjugated Hpep9.

(**a**), Chemical structures of TMR-labeled Hpep9. (**b**), HPLC-MS analysis of the purified peptide, showing the 280 nm DAD chromatogram and the isotopically resolved intact mass of the corresponding DAD peak. HRMS (ESI) analysis: calculated mass for $C_{120}H_{161}N_{31}O_{26}S$, 2484.1950; observed mass, 2484.1995.

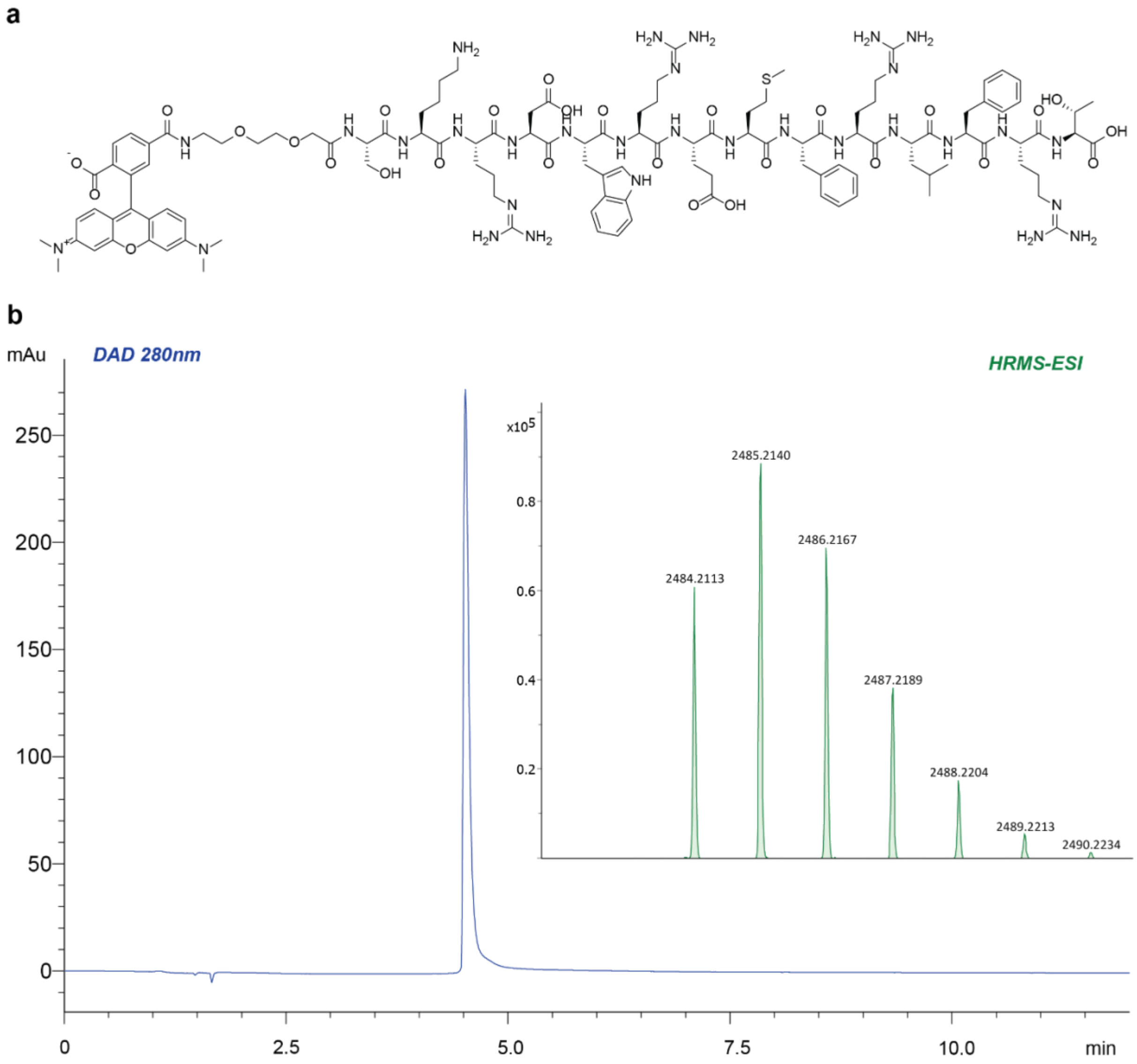

**Supplementary Fig 8.** HPLC-HRMS analysis of the TMR-conjugated Hpep11.

(**a**), Chemical structures of TMR-labeled Hpep11. (**b**), HPLC-MS analysis of the purified peptide, showing the 280 nm DAD chromatogram and the isotopically resolved intact mass of the corresponding DAD peak. HRMS (ESI) analysis: calculated mass for $C_{117}H_{165}N_{31}O_{28}S$, 2484.2161; observed mass, 2484.2113.

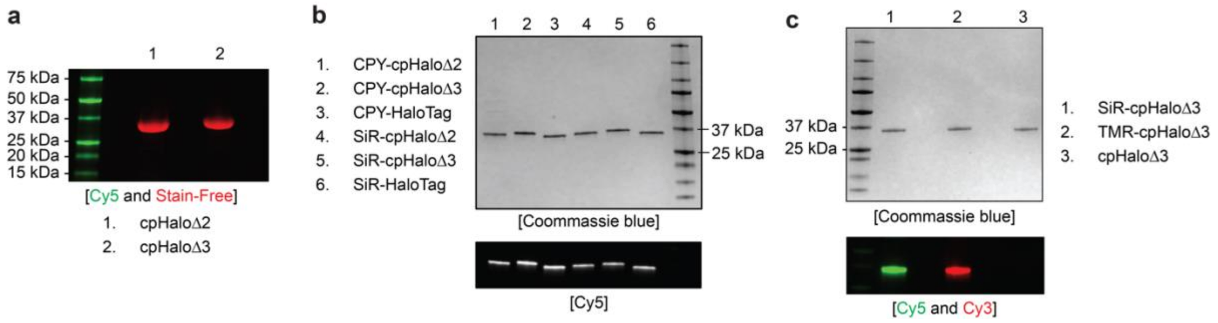

**Supplementary Fig 9.** SDS-PAGE analysis of the recombinant cpHaloΔ proteins.

The size and the labelling completeness of the cpHaloΔ proteins were verified by SDS-PAGE. The expected sizes for cpHaloΔ2 and cpHaloΔ3 is 36.4 kDa and 37.3 kDa, respectively. (**a**), Verification of the purify of cpHaloΔ proteins following His-tag purification, assessed using Stain-Free$^{TM}$ precast PAGE gels. (**b**), Verification of the fluorescently-labeled cpHaloΔ prepared for fluorescence emission scan assay (Fig.1f and Extended Data Fig. 4). (**c**), Verification of the fluorescently-labeled cpHaloΔ3 prepared for FP and BLI measurements (Extended Data Fig. 5). Fluorescence signal of SiR and CPY was collected in Cy5 channel, while TMR was collected in the Cy3 channel.

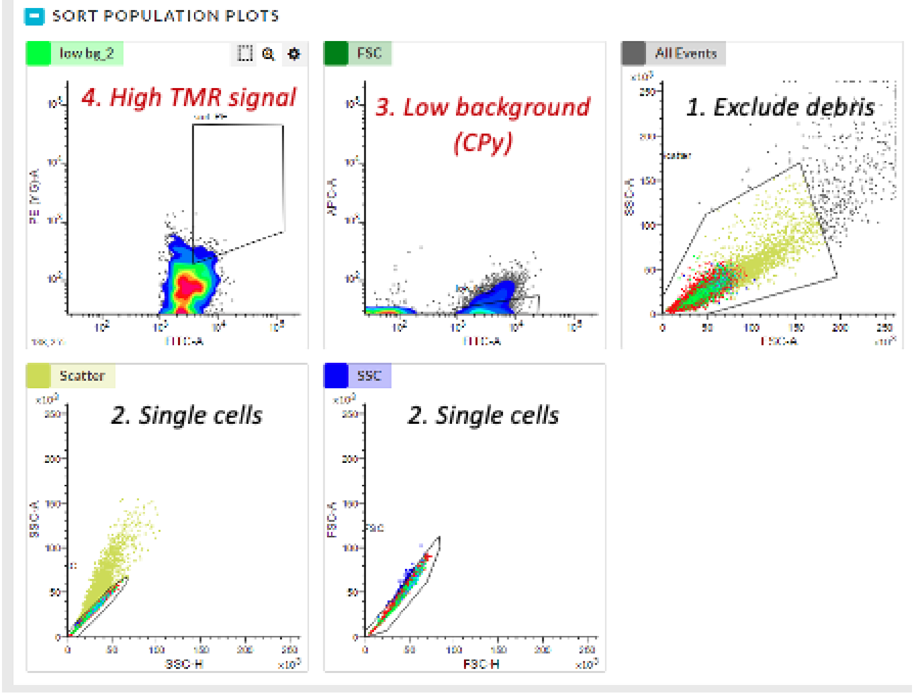

**Supplementary Fig 10.** Gating strategy for yeast library screening.

Living yeast cells were gated first using SSC-A/FSC-A, followed by singlet gating using FSC-A/FSC-H. Within the singlet population, yeast cells showing low CPY-labeling signal represented those with low residual labeling activity in absence of Hpep were gated. Within this population, the yeast cells showing high TMR signal were then sorted.

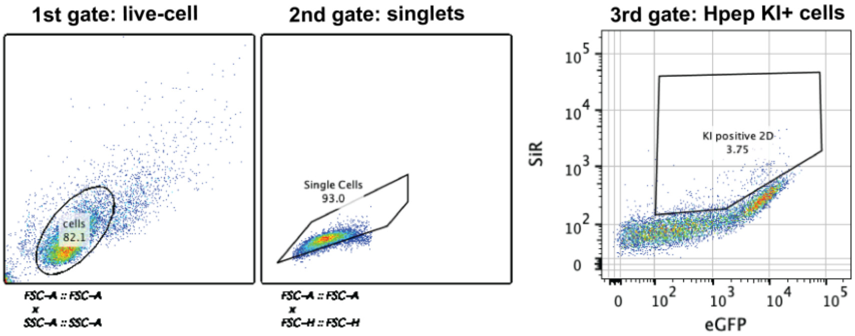

**Supplementary Fig 11.** Gating strategy to enrich Hpep11-integrated cells.

Cells were labeled with 100 nM CA-SiR for one hour. Hpep KI-positive cells co-expressing cpHaloΔ3-T2A-NLS-EGFP were enriched by FACS using the representative gating strategy.

### Supplementary Tables

**Supplementary Table 1.** $EC_{50}$ values of Hpep variants for the parental cpHaloΔ.

| Hpep | Sequence | $EC_{50}$ (µM) | 95% confidence interval |
|---|---|---|---|
| Hpep1[a] | ARETFQAFRT | 2979 | 2628 – 3417 |
| Hpep2[a] | AREMFQAFRT | 1116 | 928 – 1350 |
| Hpep3[a] | SKRDAREMFQAFRT | 149 | 128 – 176 |
| Hpep4[a] | WKEEVIKAFKLFRD | 21.0 | 18.3 -24.1 |
| Hpep5[a] | WREEVRKAFKLFRQ | 10.7 | 8.84 – 13.0 |
| Hpep6[a] | WRETFQLFRT | 2.39 | 2.08 – 2.77 |
| Hpep7[a] | WREMFRLFRT | 0.35 | 0.32 – 0.38 |
| Hpep8[a] | WKRDWREMFRLFRT | 0.12 | 0.11 – 0.14 |
| Hpep9[b] | RGWSWREMFRLFRT | 0.040 | 0.031 – 0.051 |
| Hpep10[b] | RMWTWREMFRLFRT | 0.047 | 0.030 – 0.070 |
| Hpep11[b] | SKRDWREMFRLFRT | 0.235 | 0.225 – 0.245 |

[a]The $EC_{50}$ values for Hpep1-Hpep8 are sourced from the previous publication from our group[1], and [b]the values for Hpep9-Hpep11 are reported in the PhD thesis of Jonas Wilhelm. The $EC_{50}$ characterization via CA-TMR labeling kinetics assays were done with TEVp-cleaved cpHaloΔ protein, followed by reverse-IMAC and SEC purification. For Hpeps with $EC_{50}$ values >1 µM, final assay concentrations of 500 nM cpHaloΔ protein and 100 nM CA-TMR substrate were used. For Hpeps with $EC_{50}$ values <1 µM, the assays were performed with 20 nM cpHaloΔ protein and 4 nM CA-TMR. Hpeps were titrated over a concentration range from 0 to 5 mM, depending on the $EC_{50}$ of the respective Hpep.

**Supplementary Table 2.** Affinity measurement of labeled-cpHaloΔ3 to biotin-Hpep conjugates.

| Sample | $K_d$ (M) | $K_d$ error | $k_{on}$ (1/Ms) | $k_{on}$ error | $k_{off}$ (1/s) | $k_{off}$ error | Full $R^2$ |
|---|---|---|---|---|---|---|---|
| Hpep11/ cpHaloΔ3-SiR | 4.27E-09 | 2.29E-11 | 5.00E+04 | 6.55E+01 | 2.14E-04 | 1.11E-06 | 0.9995 |
| Hpep9/ cpHaloΔ3-SiR | 1.88E-08 | 1.57E-10 | 7.89E+04 | 6.17E+02 | 1.49E-03 | 4.24E-06 | 0.996 |
| Hpep11/ cpHaloΔ3-TMR | 1.06E-08 | 2.75E-11 | 6.49E+04 | 8.97E+01 | 6.87E-04 | 1.51E-06 | 0.9993 |
| Hpep9/ cpHaloΔ3-TMR | 4.86E-09 | 6.15E-11 | 2.14E+05 | 2.30E+03 | 1.04E-03 | 1.04E-06 | 0.9899 |

The $K_d$, $k_{on}$, and $k_{off}$ error values represent the standard error of the mean (S.E.M). The full $R^2$ values were calculated from the global fit to the 1:1 binding model.

**Supplementary Table 3.** CRISPR/Cas9 KI target information.

| **Target gene name** | **Target protein_name** | **Hpep-tag location** | **crRNA_sequence (5' – 3')** | **HDR_donor_sequence**<br>lower case: homology arms; red: Hpep11; blue: linker | **RNA expression in U2OS (nTPM)** | **Localization/ subcellular structure** |
|---|---|---|---|---|---|---|
| VIM | Vimentin | N-terminus | GGACCTGGTGGACATGGCTG | **Hpep5:**cgttcgcctcttctccgggagccagtccgcgccaccgccgccgcccaggccatcgccaccctccgcagccATGAGTAAACGCGACTGGAGGGAGATGTTTAGGCTTTTTCGGACAGGTGGCGGCtccaccaggtccgtgtcctcgtcctcctaccgcaggatgttcggcggcccgggcaccgcgagccggccga | 1492.8 | Intermediate filament |
| TUBB4B | Tubulin beta-4B chain | N-terminus | CGCCGCCGCCGCCATCATGA | **Hpep5:**ctgctgctgtttgtctacttcctcctgcttccccgccgccgccgccgccatcATGAGTAAACGCGACTGGAGGGAGATGTTTAGGCTTTTTCGGACAGGTGGCGGCagggaaatcgtgcacttgcaggccgggcagtgcggcaaccaaatcggcgccaagg | 793.1 | Microtubules |
| LMNA | Prelamin-A/C | N-terminus | CCATGGAGACCCCGTCCCAG | **Hpep5:**tccttcgacccgagccccgcgccctttccgggacccctgccccgcgggcagcgctgccaacctgccggccATGAGTAAACGCGACTGGAGGGAGATGTTTAGGCTTTTTCGGACAGGTGGCGGCgagaccccgtcccagcggcgcgccacccgcagcggggcgcaggccagctccactccgctgtcgcccaccc | 255 | Nuclear speckles |
| SEC61B | Protein transport protein Sec61 subunit beta | N-terminus | GCTTGTCTCCCTCTACAGCC | **Hpep5:**gtgtctaggccggggttctggggcaggcctgccgcgctcacccgtctgtctgcttgtctccctctacagATGAGTAAACGCGACTGGAGGGAGATGTTTAGGCTTTTTCGGACAGGTGGCGGCcctggtccgacccccagtggcactaacgtgggatcctcagggcgctctcccagcaaagcagtggccgcccgggc | 248 | Endoplasmic reticulum |
| CLTA | Clathrin light chain A | N-terminus | AGCCATGGCGGGCAACTGAA | **Hpep5:**cgggcgtggtgtcggtgggtcggttggtttttgtctcaccgttggtgtccgtgccgttcagttgcccgccATGAGTAAACGCGACTGGAGGGAGATGTTTAGGCTTTTTCGGACAGGTGGCGGCgctgagctggatccgttcggcgcccctgccggcgcccctggcggtcccgcgctggggaacgga | 215 | Vesicles, endosomes, lysosomes |
| TOMM20 | Mitochondrial import receptor subunit TOM20 | C-terminus | GAGCTTGGCTGAAGATGATG | **Hpep5:**cagagaattgtaagtgctcagagcttagctgaagatgatgttgaaGGTGGCGGCAGTAAACGCGACTGGAGGGAGATGTTTAGGCTTTTTCGGACATGAgaaacaaatgtcaacataataaaatctcagtt<br>**Hpep4:**cagagaattgtaagtgctcagagctt**a**gctgaagatgatgt**t**gaaGGTGGCGGCAGAGGATGGTCATGGCGCGAAATGTTTCGGCTTTTTCGGACGTGAgaaacaaatgtcaacataataaaatctcagtt | 54.8 | Mitochondria |
| HIT2H2BE /H2BC21 | Histone H2B type 2-E | C-terminus | GGTCCCGGCAGGGACTCACT | **Hpep5:**gcccggcgagctggccaagcacgccgtgtccgagggcaccaaggcggtcaccaagtacaccagctccaagGGTGGCGGCAGTAAACGCGACTGGAGGGAGATGTTTAGGCTTTTTCGGACAtgagtccctgccgggacctggcgctcgctcgctcgagtcgccggctgcttgactccaaaggctcttttcagag | 32.2 | Nucleoplasm |

The information was sourced from the Human Protein Atlas[2]

**Supplementary Table 4.** Fluorescence lifetime of split-HaloTag pairs.

| Split-HaloTag pair | lifetime (ns) | S.E.M | N (cells) |
|---|---|---|---|
| cpHaloΔ3_cytosol | 2.285 | 0.0927 | 5 |
| cpHaloΔ3_nuclus | 2.477 | 0.0190 | 5 |
| cpHaloΔ3/Hppe8 | 3.07 | 0.017 | 5 |
| cpHaloΔ3/Hppe9 | 2.63 | 0.044 | 7 |
| cpHaloΔ3/Hppe10 | 2.22 | 0.024 | 7 |
| cpHaloΔ3/Hppe11 | 3.52 | 0.033 | 10 |

Lifetime values represent the mean measurement over the N number of cells indicated. S.E.M. = standard error of the mean.

**Supplementary Table 5.** Medium and buffers used in this study.

| Medium / Buffer | Composition |
|---|---|
| LB medium (agar) | 10 g/L tryptone, 5 g/L yeast extract, 10 g/L NaCl, pH 7.5 (15 g/L agar) |
| Activity buffer (10×) | 500 mM HEPES, 500 mM NaCl, pH 7.3 |
| Gibson assembly enzyme mix | 0.64 μL T5 exonuclease (10 U/μL), 20 μL Phusion polymerase (2 U/μL), 160 μL Taq ligase (40 U/μL), 320 μL ISO buffer (5×), 700 μL $H_2O$, aliquoted to 20 μL/reaction |
| His extract buffer | 50 mM $KH_2PO_4$, 300 mM NaCl, 5 mM imidazole, pH 8.0 |
| His washing buffer | 50 mM $KH_2PO_4$, 300 mM NaCl, 10 mM imidazole, pH 7.5 |
| His elution buffer | 50 mM $KH_2PO_4$, 300 mM NaCl, 500 mM imidazole, pH 7.5 |
| TEVp buffer | 25 mM Tris-HCl, 200 mM NaCl, 1 mM DTT, 2.5% glycerol, pH 8.0 |
| Sample buffer (4x) | 1:9 mixture of 2-mercaptoethanol and 4x Laemmli sample buffer (Bio-Rad) |
| TAE buffer (50x) | 2 M Tris, 50 mM EDTA (pH8.0), 1 M glacial acetic acid |
| YPD medium (agar) | 20 g/L tryptone, 10 g/L yeast extract, 2% glucose (15 g/L agar) |
| YPDS medium | 1:1 mixture of YPD medium and 1 M sorbitol |
| SDCAA medium (agar) | 6.7 g/L yeast nitrogen base, 5 g/L yeast synthetic drop-out medium supplement without tryptophan, 13.6 g/L $Na_2HPO_4 \cdot 12H_2O$, 8.56 g/L $NaH_2PO_4 \cdot H_2O$, 2% glucose (15 g/L agar) |
| SGCAA medium | 6.7 g/L yeast nitrogen base, 5 g/L yeast synthetic drop-out medium supplement without tryptophan, 13.6 g/L $Na_2HPO_4 \cdot 12H_2O$, 8.56 g/L $NaH_2PO_4 \cdot H_2O$, 2% galactose |
| Amino acid supplement solution | 6.7 g/L yeast nitrogen base without amino acids, 5 g/L yeast synthetic drop-out medium supplement without tryptophan |
| Electroporation buffer | 10 mM Tris-base, 0.27 M sucrose, 2.1 mM $MgCl_2$, pH 7.5 |
| DTT-Tris buffer | 0.39g dithiothreitol dissolved in 1 mL of 1 M Tris-HCl, pH 8.0 |
| LiAc-Tris buffer | 1g lithium acetate dissolved in 2 mL of 1 M Tris-HCl, pH 8.0 |
| Cell growth medium | DMEM high glucose, 10% FBS |
| Imaging medium | High-glucose phenol-red free DMEM medium, 4.5 g/L glucose, 110 mg/L pyruvate, 1x GlutaMAX™, 10% FBS |
| FACS buffer | 2% FBS in PBS (Thermo Fisher Scientific) |
| U-ExM monomer solution | 19% (w/w) sodium acrylate (SA), 10 % (w/w) AA, 0.1% (w/w) N,N'-methylenebisacrylamide (BIS) in PBS |
| 4% PFA | Diluted from 16% PFA with PBS |
| Denaturation buffer | 200 mM sodium dodecyl sulfate (SDS), 200 mM NaCl and 50 mM Tris, pH 9.0 |

**Supplementary Table 6.** Spectral properties of fluorophores used in this work.

| Fluorescent substrate | $\lambda_{abs}$ [nm] | Buffer | ε [$M^{-1}\cdot cm^{-1}$] |
|---|---|---|---|
| TMR[3] | 555 | PBS | 89,000 |
| MaP555[4] | 558 | 0.1% SDS in PBS | 142,000 |
| JFx608[5] | 608 | 10 mM HEPES, pH 7.3 | 111,000 |
| CPY[6] | 616 | 0.1% SDS in PBS | 152,000 |
| MaP618[4] | 616 | 0.1% SDS in PBS | 5,500 |
| JF635[7] | 635 | 0.1% SDS in PBS | 17,000 |
| SiR[8] | 646 | 0.1% SDS in PBS | 120,000 |
| JF646[9] | 646 | 0.1% SDS in PBS | 106,000 |
| JF669 | 674 | 0.1% SDS in PBS | 128,500 |

Extinction coefficients were extracted from literature and/or measured in-house* in the provided buffer composition.

**Supplementary Table 7.** PCR reaction recipe for cpHaloΔ library generation.

| Reagent | Volume (µL) |
|---|---|
| KOD polymerase master mix | 50 |
| $H_2O$ | 46 |
| Forward Primer 10 µM | 1.5 |
| Reverse Primer 10 µM | 1.5 |
| DNA template at 2 ngµ$L^{-1}$ | 1.0 |

**Supplementary Table 8.** PCR reaction protocol for cpHaloΔ library generation.

| Step | Temperature (°C) | Duration(s) | Cycles |
|---|---|---|---|
| Initial denaturation | 95 | 120 | 1x |
| Denaturation | 95 | 30 | |
| Primer annealing | 68 | 20 | 35x |
| DNA synthesis (elongation) | 70 | 210 | |
| Final elongation | 70 | 300 | 1x |

**Supplementary Table 9.** Digestion reaction recipe for cpHaloΔ library generation.

| Reagent | Volume (µL) |
|---|---|
| DNA (PCR product) | 60 |
| $H_2O$ | 9.5 |
| rCutSmart buffer | 8 |
| NcoI-HF or NotI-HF | 2.5 |

**Supplementary Table 10.** Ligation reaction recipe for cpHaloΔ library generation.

| Reagent | Volume (µL) |
|---|---|
| T4 DNA ligase buffer | 10 |
| $H_2O$ | 77 |
| DNA | 8 |
| T4 DNA ligase | 5 |

**Supplementary Table 11.** FACS and flow cytometry analysis.

| Fluorescent molecules | Excitation Laser (nm) | Emission Filter (nm) | Machine |
|---|---|---|---|
| eUnaG, EGFP | 488 | 530/30 | Fortessa |
| TMR | 561 | 580/15 | Fortessa |
| CPY | 640 | 670/30 | Fortessa |
| SiR | 640 | 670/30 | Fortessa |
| eUnaG, EGFP | 488 | 527/32 | Melody |
| TMR | 561 | 582/15 | Melody |
| CPY | 640 | 660/10 | Melody |
| SiR | 640 | 660/10 | Melody |

**Supplementary Table 12.** Sorting strategy for N-terminal extension cpHaloΔ library.

| Screening round | Gating | Sort count | Total events | Hpep9 concentration | Incubation time |
|---|---|---|---|---|---|
| Round 1 | counter | 17,915 | 1,706,693 | 2 μM | 30 min |
| Round 2 | counter | 20,000 | 1,403,985 | 1 μM | 30 min |
| Round 3 | counter | 10,000 | 985,765 | 500 nM | 30 min |
| Round 4 | counter | 10,000 | 884,280 | 500 nM | 15 min |
| Round 5 | negative | 20,000 | 148,213 | 250 nM | 30 min |
| Round 6 | counter | 10,000 | 915,701 | 250 nM | 30 min |

Incubation time corresponds to the time of CA-TMR (1 μM) incubation. Counter gating was performed on yeast cells labeled with CA-TMR in the presence of Hpep9, while negative gating was performed on yeast cells labeled with CA-TMR in the absence of Hpep9 as illustrated in Extended Data Fig. 1a.

**Supplementary Table 13.** Sorting strategy for C-terminal extension cpHaloΔ library.

| Screening round | Gating | Sort count | Total events | Hpep9 concentration | Incubation time |
|---|---|---|---|---|---|
| Round 1 | counter | 16,855 | 1,560,303 | 2 μM | 30 min |
| Round 2 | counter | 16,480 | 1,214,859 | 1 μM | 30 min |
| Round 3 | counter | 10,000 | 904,603 | 500 nM | 15 min |
| Round 4 | counter | 10,000 | 1,068,002 | 250 nM | 30 min |
| Round 5 | counter | 10,000 | 917,019 | 125 nM | 30 min |

**Supplementary Table 14.** Primers for library generation and NGS sample preparation.

| Library | Purpose | Forward primer sequence | Reverse primer sequence |
|---|---|---|---|
| YSD cpHaloΔ N-term | NNK library generation | **CCATGG**TAGGTTCTGGC*NNKNNKNNKNNK*GACGTCGGCCGCAAGC | **CCATGG**TACCATTAGCTGGAGCAGCC |
| YSD cpHaloΔ C-term | NNK library generation | **GCGGCCGC**TTTCTCCCAAAAGTTGG | **GCGGCCGC***MNNMNNMNNMNN*CCATTCGTCCCAGGTCGG |
| YSD cpHaloΔ N-term | NGS sample preparation | TCCCATCTATTTTCACCGCTGTTG | GCAGCGTACCCTCGATAAAAAC |
| YSD cpHaloΔ C-term | NGS sample preparation | GTTTCCACTGGGCCAAGC | GACAACGTTATCCAACAAGTTGATGTC |

Primer sequences with **restriction enzyme digestion site**, degenerate codons, and annealing sequences specified.

**Supplementary Table 15.** Stable cell lines generated in this study.

| Cell type | Plasmid 1 | Stable integration method | Plasmid 2 | Stable integration method | Figure |
|---|---|---|---|---|---|
| U2OS | pcDNA5/FRT/TO-EGFP-GSG-cpHaloΔ2 | Flp-IN | - | - | Extended Data Fig. 3 |
| U2OS | pcDNA5/FRT/TO-EGFP-GSG-cpHaloΔ3 | Flp-IN | - | - | Fig. 3f-g;<br>Extended Data Fig. 3 |
| U2OS | pcDNA5/FRT/TO-H2B-SNAPf-Hpep11 | Flp-IN | - | - | Fig. 2c;<br>Extended Data Fig. 6d |
| U2OS | pcDNA5/FRT/TO-TOMM20-SNAPf-Hpep11 | Flp-IN | - | - | Fig. 2c;<br>Extended Data Fig. 6d |
| U2OS | pAAVS1-P-TO-EGFP-GSG-cpHaloΔ3 | AAVS1 safe harbor | pcDNA5/FRT/TO-TOMM20-SNAPf-Hpep11 | Flp-IN | Fig. 2d; Fig. 3f, 3g; Fig. 4a;<br>Extended Data Fig. 6c |
| U2OS | GGG-Hpep or Hpep-GGG | CRISPR/Cas9 KI | pcDNA5/FRT/TO- cpHaloΔ3-T2A-NLS-EGFP | Flp-IN | Fig. 3; Fig. 4; Fig. 5b;<br>Extended Data Fig. 8a-c;<br>Extended Data Fig. 8e |
| U2OS | GGG-HaloTag | CRISPR/Cas9 KI | - | - | Fig. 3 |
| U2OS | HaloTag-GGG | CRISPR/Cas9 KI | - | - | Fig. 4d |
| U2OS | pcDNA5/FRT/TO-cpHaloΔ3-T2A-NLS-EGFP | Flp-IN | - | - | Fig. 3c-d;<br>Extended Data Fig. 8e |
| 293T | pAAVS1-P-TO_cpHdel75-T2A-NLS-EGFP | AAVS1 safe harbor | GGG-Hpep11 | CRISPR/Cas9 KI | Extended Data Fig. 8d |
| 293T | GGG-HaloTag | CRISPR/Cas9 KI | - | - | Extended Data Fig. 8d |

**Supplementary Table 16.** U2OS transient transfection and associated experimental figures.

| 1st expression cassette (stable expression) | 2nd expression cassette (stable expression) | 3rd expression cassette (plasmid transient transfection) | Figure | Annotation |
|---|---|---|---|---|
| EGFP-GSG-cpHaloΔ3 | - | H2B-SNAPf-Hpep9 | Fig. 2a | |
| EGFP-GSG-cpHaloΔ3 | - | TOMM20-SNAPf-Hpep9 | Fig. 2a | |
| EGFP-GSG-cpHaloΔ3 | - | Hpep9-SNAPf-LamB1 | Fig. 2a | |
| EGFP-GSG-cpHaloΔ3 | - | H2B-SNAPf-Hpep11 | Fig. 2b | |
| EGFP-GSG-cpHaloΔ3 | - | TOMM20-SNAPf-Hpep11 | Fig. 2b | |
| EGFP-GSG-cpHaloΔ3 | - | Hpep11-SNAPf-LamB1 | Fig. 2b | |
| - | - | TOMM20-SNAPf-Hpep11 | Extended Data Fig. 6a | |
| - | - | EGFP-GSG-cpHaloΔ3 | Extended Data Fig. 6a | |
| - | - | Hpep11-SNAPf-LamB1 | Extended Data Fig. 6b | |
| EGFP-GSG-cpHaloΔ3 | - | Hpep11-SNAPf-LamB1 | Extended Data Fig. 6b | |
| cpHaloΔ3-T2A-NLS-EGFP | - | Hpep11-SNAPf-LamB1 | Extended Data Fig. 6b | |
| cpHaloΔ3-T2A-NLS-EGFP | TOMM20-SNAPf-Hpep11 | H2B-SNAPf-Hpep8 | Fig. 5d | |
| cpHaloΔ3-T2A-NLS-EGFP | TOMM20-SNAPf-Hpep11 | H2B-SNAPf-Hpep9 | Fig. 5d | |
| cpHaloΔ3-T2A-NLS-EGFP | TOMM20-SNAPf-Hpep11 | H2B-SNAPf-Hpep10 | Fig. 5d | |
| - | - | cpHaloΔ3-T2A-EGFP-P30-SNAPf-NLS3x | Extended Data Fig. 7b | Control |
| - | - | cpHaloΔ3-T2A-EGFP-Hpep11[loopC_gsgx2]-P30-SNAPf-NLS3x | Extended Data Fig. 7b | LoopC |
| - | - | EGFP-cpHaloTag[Hpep11+cpHaloΔ3][loopC_gsgx2]-P30-SNAPf-NLS3x | Extended Data Fig. 7b | LoopC |
| - | - | EGFP-HaloTag7[loopC_gsgx2]-P30-SNAPf-NLS3x | Extended Data Fig. 7b | LoopC |
| - | - | cpHaloΔ3-T2A-EGFP-gsg-Hpep11-P30-SNAPf-NLS3x | Extended Data Fig. 7b | C-terminus |
| - | - | EGFP-gsg-cpHaloTag[Hpep11+cpHaloΔ3]-P30-SNAPf-NLS3x | Extended Data Fig. 7b | C-terminus |
| - | - | EGFP-gsg-HaloTag7-P30-SNAPf-NLS3x | Extended Data Fig. 7b | C-terminus |
| - | - | cpHaloΔ3-T2A-gsg-Hpep11-gsg-EGFP-P30-SNAPf-NLS3x | Extended Data Fig. 7b | N-terminus |
| - | - | cpHaloTag[Hpep11+cpHaloΔ3]-gsg-EGFP-P30-SNAPf-NLS3x | Extended Data Fig. 7b | N-terminus |
| - | - | HaloTag7-gsg-EGFP-P30-SNAPf-NLS3x | Extended Data Fig. 7b | N-terminus |

All the above-listed mammalian expression plasmids were constructed using the pcDNA5/FRT/TO vector.

**Supplementary Table 17.** Confocal image acquisition parameters.

| Figure | Hpep tagging target/construct (s) | Ligands for labeling (s) | Objectives | Excitation [nm] (laser power/ch.) | Emission [nm] | Pixel dwell time [µs] | Pinhole [Airy units, mAU] | Pixel size [nm] | Size [pixels] | Comments |
|---|---|---|---|---|---|---|---|---|---|---|
| 2a | H2B-STf-Hpep9 | CP-TMR, CA-SiR | 40x/1.10 water | 485 (0.35% EGFP), 550 (0.3% TMR), 640 (0.1% SiR) | 495-530, 560-600, 650-720 | 1.75 | 999.46 | 90 | 928x928 | 2 line-average |
| 2a | TOM20-STf-Hpep9 | CP-TMR, CA-SiR | 40x/1.10 water | 485 (1.5% EGFP), 550 (1% TMR), 640 (0.5% SiR) | 495-530, 560-600, 650-720 | 1.3875 | 999.46 | 90 | 1168x1168 | 2 line-average |
| 2a | TOM20-STf-Hpep9 (zoom-in) | CP-TMR, CA-SiR | 40x/1.10 water | 485 (3% EGFP), 550 (1% TMR), 640 (0.5% SiR) | 495-530, 560-600, 650-720 | 14.5 | 999.46 | 87 | 112x112 | 2 line-average |
| 2a | Hpep9-STf-LamB1 | CP-TMR, CA-SiR | 40x/1.10 water | 485 (0.3% EGFP), 555 (0.2% TMR), 640 (2% SiR) | 495-530, 565-620, 650-720 | 0.5 | 999.46 | 90 | 3248x3248 | 2 line-average |
| 2b | TOM20-STf-Hpep11 | CP-TMR, CA-SiR | 40x/1.10 water | 485 (3% EGFP), 555 (1% TMR), 640 (1% SiR) | 495-530, 565-620, 650-720 | 0.7875 | 999.46 | 41 | 2048x2048 | 2 line-average |
| 2b | H2B-STf-Hpep11 | CP-TMR, CA-SiR | 40x/1.10 water | 485 (10% EGFP), 555 (1% TMR), 640 (1% SiR) | 495-530, 565-620, 650-720 | 0.7875 | 999.46 | 57 | 2048x2048 | 2 line-average |
| 2b | Hpep11-STf-LamB1 | CP-TMR, CA-SiR | 40x/1.10 water | 485 (4.5% EGFP), 555 (2% TMR), 640 (15% SiR) | 495-530, 565-620, 650-720 | 0.7875 | 999.46 | 57 | 2048x2048 | 2 line-average |
| 2c | H2B-STf-Hpep11 | cpHaloΔ3, CA-SiR, CP-TMR | 40x/1.10 water | 555 (1% TMR) 645 (2% SiR) | 565-600 655-720 | 0.85 | 999.46 | 102 | 1896x1896 | 2 line-average |

| Figure | Hpep tagging target/construct (s) | Ligands for labeling (s) | Objectives | Excitation [nm] (laser power/ch.) | Emission [nm] | Pixel dwell time [µs] | Pinhole [Airy units, mAU] | Pixel size [nm] | Size [pixels] | Comments |
|---|---|---|---|---|---|---|---|---|---|---|
| 2c | TOM20-STf-Hpep11 | cpHaloΔ3, CA-SiR, CP-TMR | 40x/1.10 water | 555 (6% TMR)<br>645 (25% SiR) | 565-600<br>655-720 | 1.575 | 999.46 | 142 | 1024x1024 | 2 line-average |
| 2c | U2OS blank cells | cpHaloΔ3, CA-SiR, CP-TMR | 40x/1.10 water | 555 (6% TMR)<br>645 (25% SiR) | 565-600<br>655-720 | 0.85 | 999.46 | 102 | 1896x1896 | 2 line-average |
| 2d | TOM20-STf-Hpep11 | CA-MaP555, | 40x/1.10 water | 555 (100%) | 565-620 | 1.575 | 999.46 | 142 | 1024x1024 | 2 line-average |
| 2d | TOM20-STf-Hpep11 | CA-TMR | 40x/1.10 water | 555 (21.25%) | 565-620 | 1.575 | 999.46 | 142 | 1024x1024 | 2 line-average |
| 2d | TOM20-STf-Hpep11 | CA-CPY | 40x/1.10 water | 610 (7.27%) | 620-700 | 3.8375 | 999.46 | 142 | 1024x1024 | 2 line-average |
| 3b | Hpep11-Sec61B | CA-CPY | 40x/1.10 water | 485 (1% EGFP),<br>615 (3.5% CPY) | 495-532,<br>625-700 | 1.4625 | 999.46 | 90 | 1104x1104 | 2 line-average |
| 3b | Hpep11-VIM | CA-CPY | 40x/1.10 water | 485 (0.3% EGFP),<br>615 (2% CPY) | 495-532,<br>625-700 | 1.45 | 999.46 | 89 | 1120x1120 | 2 line-average |
| 3b | Hpep11-CLTA | CA-CPY | 40x/1.10 water | 485 (0.6% EGFP),<br>615 (1.5% CPY) | 495-532,<br>625-700 | 1.4875 | 999.46 | 89 | 1088x1088 | 2 line-average |
| 3b | Hpep11-LMNA | CA-CPY | 40x/1.10 water | 485 (1% EGFP),<br>615 (1.8% CPY) | 495-532,<br>625-700 | 1.2375 | 999.46 | 89 | 1304x1304 | 2 line-average |
| 3b | HIST2H2BE-Hpep11 | CA-CPY | 40x/1.10 water | 485 (0.7% EGFP),<br>615 (0.3% CPY) | 495-532,<br>625-700 | 0.9875 | 999.46 | 89 | 1640x1640 | 2 line-average |
| 3b | TOMM20-Hpep11 | CA-CPY | 40x/1.10 water | 485 (2% EGFP),<br>615 (2% CPY) | 495-532,<br>625-700 | 1.2125 | 999.46 | 89 | 1328x1328 | 2 line-average |
| Extended Data Fig.8a | Hpep11-LMNA | CA-SiR | 40x/1.10 water | 485 (1% EGFP),<br>640 (1.5% SiR) | 495-532,<br>650-720 | 1.7 | 999.46 | 89 | 952x952 | 2 line-average |

| Figure | Hpep tagging target/construct (s) | Ligands for labeling (s) | Objectives | Excitation [nm] (laser power/ch.) | Emission [nm] | Pixel dwell time [µs] | Pinhole [Airy units, mAU] | Pixel size [nm] | Size [pixels] | Comments |
|---|---|---|---|---|---|---|---|---|---|---|
| Extended Data Fig.8a | Hpep11-CLTA | CA-SiR | 40x/1.10 water | 485 (0.3% EGFP), 640 (2% SiR) | 495-532, 650-720 | 1.3 | 999.46 | 89 | 1248x1248 | 2 line-average |
| Extended Data Fig.8a | Hpep11-Sec61B | CA-SiR | 40x/1.10 water | 485 (0.6% EGFP), 640 (3% SiR) | 495-532, 650-720 | 1.2375 | 999.46 | 89 | 1304x1304 | 2 line-average |
| Extended Data Fig.8a | HIST2H2BE-Hpep11 | CA-SiR | 40x/1.10 water | 485 (1% EGFP), 640 (0.3% SiR) | 495-532, 650-720 | 1.2125 | 999.46 | 89 | 1336x1336 | 2 line-average |
| Extended Data Fig.8a | TOMM20-Hpep11 | CA-SiR | 40x/1.10 water | 485 (0.6% EGFP), 640 (3% SiR) | 495-532, 650-720 | 1.3 | 999.46 | 90 | 1248x1248 | 2 line-average |
| 3d | TOMM20-Hpep9/Hpep11/HT7 | CA-SiR | 40x/1.10 water | 485 (1.8% EGFP), 640 (10% SiR) | 495-530, 650-720 | 1.4375 | 999.46 | 89 | 1128x1128 | |
| Extended Data Fig.8c | HaloTag-NOP10 | CA-SiR | 40x/1.10 water | 488 (2% EGFP), 640 (3% SiR) | 498-520, 650-720 | 1 | 999.46 | 90 | 1616x1616 | |
| Extended Data Fig.8c | Hpep11-NOP10 | CA-SiR | 40x/1.10 water | 488 (0.5% EGFP), 640 (1% SiR) | 498-520, 650-720 | 1 | 999.46 | 90 | 1616x1616 | |
| 3e and Extended Data Fig.8d | HIST2H2BE-Hpep11 | CA-TMR, CA-MaP555, CA-JFx608, CA-CPY, CA-MaP618, CA-JF635, CA-SiR, CA-JF646, CA- JF669 | 40x/1.10 water | 488 (1% EGFP), 555 (1% TMR or MaP555), 608 (0.1% JFx608), 610 (0.1% CPY), 618 (2% MaP618), 635 (2% JF635), 640 (0.2% SiR), 646 (0.2% JF646), 669 (0.6% JF669) | 498-530, 565-620, 618-720, 620-720, 628-720, 645-720, 650-720, 656-720, 679-720 | 0.7875 | 999.46 | 142 | 2048x2048 | |

| Figure | Hpep tagging target/construct (s) | Ligands for labeling (s) | Objectives | Excitation [nm] (laser power/ch.) | Emission [nm] | Pixel dwell time [µs] | Pinhole [Airy units, mAU] | Pixel size [nm] | Size [pixels] | Comments |
|---|---|---|---|---|---|---|---|---|---|---|
| 3e and Extended Data Fig.8e | Hpep11-LMNA | CA-TMR,<br>CA-MaP555,<br>CA-JFx608,<br>CA-CPY,<br>CA-MaP618,<br>CA-JF635,<br>CA-SiR,<br>CA-JF646,<br>CA- JF669 | 40x/1.10 water | 488 (1% EGFP),<br>555 (20% TMR or 50% MaP555),<br>608 (0.8% JFx608),<br>610 (0.5% CPY),<br>618 (80% MaP618),<br>635 (80% JF635),<br>640 (8% SiR),<br>646 (15% JF646),<br>669 (50% JF669) | 498-530,<br>565-620,<br>618-720,<br>620-720,<br>628-720,<br>645-720,<br>650-720,<br>656-720,<br>679-720 | 0.7875 | 999.46 | 142 | 2048x2048 | |
| 5c | TOMM20-Hpep11 | CA-SiR | 40x/1.10 water | 640 (60% SiR) | 650-720 | 5.125 | 999.37 | 118 | 768x768 | CRISPR cells<br>80 MHz, SP8 |
| 5c | Hpep11-STf-LamB1 | CA-SiR | 40x/1.10 water | 640 (20% SiR) | 650-720 | 4.2 | 999.37 | 118 | 936x936 | 80 MHz, SP8 |
| 5d | H2B-STf-Hpep10 | CA-SiR | 40x/1.10 water | 640 (3% SiR) | 650-720 | 3.4625 | 999.37 | 118 | 616x616 | 80 MHz, SP8 |
| 5d | H2B-STf-Hpep8 | CA-SiR | 40x/1.10 water | 640 (3% SiR) | 650-720 | 3.425 | 999.37 | 118 | 624x624 | 80 MHz, SP8 |
| 5d | H2B-STf-Hpep9 | CA-SiR | 40x/1.10 water | 640 (1% SiR) | 650-720 | 3.6125 | 999.37 | 117 | 592x592 | 80 MHz, SP8 |
| Extended Data Fig.6a | | CP-TMR,<br>CA-SiR | 40x/1.10 water | 485 (3%),<br>555 (2% TMR),<br>640 (0.2% SiR) | 495-530,<br>565-620,<br>650-720 | 0.7875 | 999.46 | 71 | 2048x2048 | |
| Extended Data Fig.6b | | CP-TMR,<br>CA-SiR | 40x/1.10 water | 485 (1.5%),<br>555 (1% TMR),<br>640 (4.5% SiR) | 495-530,<br>565-620,<br>650-720 | 1.5375 | 999.46 | 92 | 1056x1056 | |

| Figure | Hpep tagging target/construct (s) | Ligands for labeling (s) | Objectives | Excitation [nm] (laser power/ch.) | Emission [nm] | Pixel dwell time [µs] | Pinhole [Airy units, mAU] | Pixel size [nm] | Size [pixels] | Comments |
|---|---|---|---|---|---|---|---|---|---|---|
| Extended Data Fig.6c | | CP-TMR, CA-CPY | 40x/1.10 water | 485 (10% EGFP), 550 (1.6% TMR), 600 (1.5% SiR) | 495-530, 565-595, 610-700 | 1.575 | 999.46 | 284 | 1024x1024 | Live-cell |
| Extended Data Fig.6c | | CP-TMR, CA-CPY | 40x/1.10 water | 485 (1.5% EGFP), 550 (3.5% TMR), 600 (0.7% SiR) | 495-530, 565-595, 610-700 | 1.575 | 999.46 | 142 | 1024x1024 | Fixed cell |
| Extended Data Fig.6d | H2B-STf-Hpep11 | cpHaloΔ3, CA-CPY, CP-TMR | 40x/1.10 water | 555 (0.5% TMR), 615 (0.5% CPY) | 565-600, 625-720 | 1.575 | 999.46 | 189 | 1024x1024 | |
| Extended Data Fig.6d | TOM20-STf-Hpep11 | cpHaloΔ3, CA-CPY, CP-TMR | 40x/1.10 water | 555 (6% TMR), 615 (15% CPY) | 565-600, 625-720 | 1.575 | 999.46 | 189 | 1024x1024 | |
| Extended Data Fig.6d | U2OS blank cells | cpHaloΔ3, CA-CPY, CP-TMR | 40x/1.10 water | 555 (6% TMR), 615 (25% CPY) | 565-600, 625-720 | 0.85 | 999.46 | 102 | 1896x1896 | |
| Extended Data Fig.7c | | CP-TMR, CA-SiR | 40x/1.10 water | 485 (1.5% EGFP), 550 (0.3% TMR), | 495-530, 560-620, | 0.7875 | 999.46 | 142 | 2048x2048 | |

All the above-mentioned image acquisitions were performed on Stellaris 5 confocal microscope unless otherwise stated.

**Supplementary Table 18.** STED image acquisition parameters.

| Figure | Target | Cell-lines | Imaging | Ex (%) | STED (%) | Pixel dwell time [µs] | Pixel size [nm] | Size [µm] | Emission [nm] | Comments |
|---|---|---|---|---|---|---|---|---|---|---|
| 4a (upper) | TOM20-STf-Hpep11 + EGFP-GSG-cpHaloΔ3 | KI | CLSM<br>STED | 10<br>13 | -<br>10 | 15 | 30 | 10 x 10 | 650-757 | 3 line accu. |
| 4a (lower) | TOM20-GGG-Hpep11 + cpHaloΔ3-T2A-NLS-EGFP | FTR | CLSM<br>STED | 10<br>13 | -<br>10 | 15 | 30 | 10 x 10 | 650-757 | 3 line accu. |
| 4b (upper) | TOM20-HT7 | KI | CLSM<br>STED | 14<br>18 | -<br>10 | 15 | 30 | 10 x 10 | 650-757 | 3 line accu. |
| 4b (lower) | TOM20-GGG-Hpep11 + cpHaloΔ3-T2A-NLS-EGFP | KI | CLSM<br>STED | 14<br>18 | -<br>10 | 15 | 30 | 10 x 10 | 650-757 | 3 line accu. |
| 4c | Hpep11-GGG-CLTA + cpHaloΔ3-T2A-NLS-EGFP | KI | CLSM ov<br>CLSM z<br>STED | 5<br>5<br>15 | -<br>-<br>10 | 15 | 80<br>50<br>30 | 50 x 50<br>10 x 10<br>10 x 10 | 650-757 | 2 line accu. |
| 4d (left) | Hpep11-GGG-TUBB4B + cpHaloΔ3-T2A-NLS-EGFP | KI | CLSM ov<br>CLSM z<br>STED | 2<br>3<br>5 | -<br>-<br>20 | 10 | 80<br>50<br>30 | 50 x 50<br>10 x 10<br>10 x 10 | 650-757 | 3 line accu. |
| 4d (right) | HaloTag7-GGG-TUBB4B | KI | CLSM ov<br>CLSM z<br>STED | 2<br>3<br>5 | -<br>-<br>20 | 10 | 80<br>50<br>30 | 50 x 50<br>10 x 10<br>10 x 10 | 650-757 | 3 line accu. |

Excitation line: 640 nm; STED line: 775 nm. KI – Knock-in, FTR – overexpression via Flp-In T-REx system. ov – overview, z – zoom.

### Protein sequences

#### Expression in *E.coli*

>cpHaloΔ

MHHHHHHHHHHENLYFQGDVGRKLIIDQNVFIEGTLPMGVVRPLTEVEMDHYREPFLNPVDREPLWRFPNELPIAGEPANIV
ALVEEYMDWLHQSPVPKLLFWGTPGVLIPPAEAARLAKSLPNCKAVDIGPGLNLLQEDNPDLIGSEIARWLSTLEIGGTGGSGGT
GGSGGSIGTGFPFDPHYVEVLGERMHYVDVGPRDGTPVLFLHGNPTSSYVWRNIIPHVAPTHRCIAPDLIGMGKSDKPDLGYFFD
DHVRFMDAFIEALGLEEVVLVIHDWGSALGFHWAKRNPERVKGIAFMEFIRPIPTWDEW*

His-tag – TEVp site – cpHaloΔ

>cpHaloΔ2

MHHHHHHHHHHENLYFQGDVGRKLIIDQNVFIEGTLPMGVVRPLTEEEMDHYREPFLNPKDREPLWRFPNELPIAGEPANIV
ALVEEYMDWLHQSPVPKLLFWGTPGVLIPPAEAARLAKSLPNCKAVDIGPGLNLLQEDNPDLIGSEIARWLSTLEIKSKYDRDQI
LKIIAELEKKTGGSIGTGFPFDPHYVEVLGSRMHYVDVGPRDGTPVLFLHGNPTSSYVWRNIIPHVAPTHRCIAPDLIGMGKSDKP
DLGYFFDDHVRFMDAFIEALGLEEVVLVIHDWGSALGFHWAKRHPERVKGIAFMEFIRPIPTWDEW*

His-tag – TEVp site – cpHaloΔ2

>cpHaloΔ3

MHHHHHHHHHHENLYFQGEKKGDVGRKLIIDQNVFIEGTLPMGVVRPLTEEEMDHYREPFLNPKDREPLWRFPNELPIAGEP
ANIVALVEEYMDWLHQSPVPKLLFWGTPGVLIPPAEAARLAKSLPNCKAVDIGPGLNLLQEDNPDLIGSEIARWLSTLEIKSKYD
RDQILKIIAELEKKTGGSIGTGFPFDPHYVEVLGSRMHYVDVGPRDGTPVLFLHGNPTSSYVWRNIIPHVAPTHRCIAPDLIGMGK
SDKPDLGYFFDDHVRFMDAFIEALGLEEVVLVIHDWGSALGFHWAKRHPERVKGIAFMEFIRPIPTWDEWGDVE*

His-tag – TEVp site – cpHaloΔ3

#### Expression on yeast surface

>pJYDNg-cpHaloΔ_N-term extension

MRFPSIFTAVVFAASSALAAPANGTMVGSGXXXXDVGRKLIIDQNVFIEGTLPMGVVRPLTEVEMDHYREPFLNPVDREPLWRF
PNELPIAGEPANIVALVEEYMDWLHQSPVPKLLFWGTPGVLIPPAEAARLAKSLPNCKAVDIGPGLNLLQEDNPDLIGSEIARWL
STLEIGGTGGSGGTGGSGGSIGTGFPFDPHYVEVLGERMHYVDVGPRDGTPVLFLHGNPTSSYVWRNIIPHVAPTHRCIAPDLIG
MGKSDKPDLGYFFDDHVRFMDAFIEALGLEEVVLVIHDWGSALGFHWAKRNPERVKGIAFMEFIRPIPTWDEWAAAFSQKLDI
NLLDNVVNSSYHGEGVSGGSAQELTTICEQIPSPTLESTPYSLSTTTILANGKAMQGVFEYYKSVTFVSNCGSHPSTTSKGSPINTQ
YVFKDNSSTIEGRYPYDVPDYALQASGGGGSGGGGSGGGGSASHEQKLISEEDLMLEKFVGTWKIESSENFGEYLKAIGAPKELAD
AGDATTPVLYISQKDGDKMTVKIENGPPTFLDTQVSFKLGEEFDEFPSDRRKGVKSVVNLSGEKLVYVQKWDGKETTYVREIKD
GKLVVTLTMGDVVAVRSYRRASE**

Leader sequence appS4 – cpHaloΔ_Lib – Aga2p – Factor Xa site – HA-tag – Myc-tag – Linker – eUnaG2

>pJYDNg-cpHaloΔ_C-term extension

MRFPSIFTAVVFAASSALAAPANGTMVDVGRKLIIDQNVFIEGTLPMGVVRPLTEVEMDHYREPFLNPVDREPLWRFPNELPIA
GEPANIVALVEEYMDWLHQSPVPKLLFWGTPGVLIPPAEAARLAKSLPNCKAVDIGPGLNLLQEDNPDLIGSEIARWLSTLEIGG
TGGSGGTGGSGGSIGTGFPFDPHYVEVLGERMHYVDVGPRDGTPVLFLHGNPTSSYVWRNIIPHVAPTHRCIAPDLIGMGKSDKP
DLGYFFDDHVRFMDAFIEALGLEEVVLVIHDWGSALGFHWAKRNPERVKGIAFMEFIRPIPTWDEWXXXXAAAFSQKLDINLL
DNVVNSSYHGEGVSGGSAQELTTICEQIPSPTLESTPYSLSTTTILANGKAMQGVFEYYKSVTFVSNCGSHPSTTSKGSPINTQYVF
KDNSSTIEGRYPYDVPDYALQASGGGGSGGGGSGGGGSASHEQKLISEEDLMLEKFVGTWKIESSENFGEYLKAIGAPKELADAG
DATTPVLYISQKDGDKMTVKIENGPPTFLDTQVSFKLGEEFDEFPSDRRKGVKSVVNLSGEKLVYVQKWDGKETTYVREIKDGK
LVVTLTMGDVVAVRSYRRASE**

Leader sequence appS4 – cpHaloΔ_Lib – Aga2p – Factor Xa site – HA-tag – Myc-tag – Linker – eUnaG2

#### Expression in mammalian cells

> cpHaloΔ

DVGRKLIIDQNVFIEGTLPMGVVRPLTEVEMDHYREPFLNPVDREPLWRFPNELPIAGEPANIVALVEEYMDWLHQSPVPKLLF
WGTPGVLIPPAEAARLAKSLPNCKAVDIGPGLNLLQEDNPDLIGSEIARWLSTLEIGGTGGSGGTGGSGGSIGTGFPFDPHYVEVL
GERMHYVDVGPRDGTPVLFLHGNPTSSYVWRNIIPHVAPTHRCIAPDLIGMGKSDKPDLGYFFDDHVRFMDAFIEALGLEEVVL
VIHDWGSALGFHWAKRNPERVKGIAFMEFIRPIPTWDEW

> cpHaloΔ2

DVGRKLIIDQNVFIEGTLPMGVVRPLTEEEMDHYREPFLNPKDREPLWRFPNELPIAGEPANIVALVEEYMDWLHQSPVPKLLF
WGTPGVLIPPAEAARLAKSLPNCKAVDIGPGLNLLQEDNPDLIGSEIARWLSTLEIKSKYDRDQILKIIAELEKKTGGSIGTGFPFD
PHYVEVLGSRMHYVDVGPRDGTPVLFLHGNPTSSYVWRNIIPHVAPTHRCIAPDLIGMGKSDKPDLGYFFDDHVRFMDAFIEAL
GLEEVVLVIHDWGSALGFHWAKRHPERVKGIAFMEFIRPIPTWDEW

> cpHaloΔ3

EKKGDVGRKLIIDQNVFIEGTLPMGVVRPLTEEEMDHYREPFLNPKDREPLWRFPNELPIAGEPANIVALVEEYMDWLHQSPVP
KLLFWGTPGVLIPPAEAARLAKSLPNCKAVDIGPGLNLLQEDNPDLIGSEIARWLSTLEIKSKYDRDQILKIIAELEKKTGGSIGTG
FPFDPHYVEVLGSRMHYVDVGPRDGTPVLFLHGNPTSSYVWRNIIPHVAPTHRCIAPDLIGMGKSDKPDLGYFFDDHVRFMDA
FIEALGLEEVVLVIHDWGSALGFHWAKRHPERVKGIAFMEFIRPIPTWDEWGDVE
N/C-terminal extensions – cpHaloΔ2

> HaloTag

GSEIGTGFPFDPHYVEVLGERMHYVDVGPRDGTPVLFLHGNPTSSYVWRNIIPHVAPTHRCIAPDLIGMGKSDKPDLGYFFDDH
VRFMDAFIEALGLEEVVLVIHDWGSALGFHWAKRNPERVKGIAFMEFIRPIPTWDEWPEFARETFQAFRTTDVGRKLIIDQNVF
IEGTLPMGVVRPLTEVEMDHYREPFLNPVDREPLWRFPNELPIAGEPANIVALVEEYMDWLHQSPVPKLLFWGTPGVLIPPAE
AARLAKSLPNCKAVDIGPGLNLLQEDNPDLIGSEIARWLSTLEISG*

>pcDNA5/FRT/TO-cpHaloΔ-T2A-NLS-EGFP

M[cpHaloΔ]GSGATNFSLLKQAGDVEENPGPSRMAPKKKRKVMVSKGEELFTGVVPILVELDGDVNGHKFSVSGEGEGDATYGK
LTLKFICTTGKLPVPWPTLVTTLTYGVQCFSRYPDHMKQHDFFKSAMPEGYVQERTIFFKDDGNYKTRAEVKFEGDTLVNRIEL
KGIDFKEDGNILGHKLEYNYNSHNVYIMADKQKNGIKVNFKIRHNIEDGSVQLADHYQQNTPIGDGPVLLPDNHYLSTQSALSK
DPNEKRDHMVLLEFVTAAGITLGMDELYK**
cpHaloΔ – T2A (2A self-cleaving sequence) – NLS (nuclear localization signal or sequence) – Linker – EGFP

>pcDNA5/FRT/TO-EGFP-cpHaloΔ

MVSKGEELFTGVVPILVELDGDVNGHKFSVSGEGEGDATYGKLTLKFICTTGKLPVPWPTLVTTLTYGVQCFSRYPDHMKQHDF
FKSAMPEGYVQERTIFFKDDGNYKTRAEVKFEGDTLVNRIELKGIDFKEDGNILGHKLEYNYNSHNVYIMADKQKNGIKVNFKI
RHNIEDGSVQLADHYQQNTPIGDGPVLLPDNHYLSTQSALSKDPNEKRDHMVLLEFVTAAGITLGMDELYKGSGGTGGSG[cpH
aloΔ]*
EGFP – Linker – cpHaloΔ

>pcDNA5/FRT/TO-H2B-SNAPf-Hpep

MPEPAKSAPAPKKGSKKAVTKAQKKGGKKRKRSRKESYSIYVYKVLKQVHPDTGISSKAMGIMNSFVNDIFERIAGEASRLAHY
NKRSTITSREIQTAVRLLLPGELAKHAVSEGTKAITKYTSAKGGDKDCEMKRTTLDSPLGKLELSGCEQGLHRIIFLGKGTSAADA
VEVPAPAAVLGGPEPLMQATAWLNAYFHQPEAIEEFPVPALHHPVFQQESFTRQVLWKLLKVVKFGEVISYSHLAALAGNPAA
TAAVKTALSGNPVPILIPCHRVVQGDLDVGGYEGGLAVKEWLLAHEGHRLGKPGLGSG[Hpep]*
H2B – Linker – SNAPf – Hpep

>pcDNA5/FRT/TO-LamB1-SNAPf-Hpep

MATATPVPPRMGSRAGGPTTPLSPTRLSRLQEKEELRELNDRLAVYIDKVRSLETENSALQLQVTEREEVRGRELTGLKALYET
ELADARRALDDTARERAKLQIELGKCKAEHDQLLLNYAKKESDLNGAQIKLREYEAALNSKDAALATALGDKKSLEGDLEDLKD
QIAQLEASLAAAKKQLADETLLKVDLENRCQSLTEDLEFRKSMYEEEINETRRKHETRLVEVDSGRQIEYEYKLAQALHEMREQ
HDAQVRLYKEELEQTYHAKLENARLSSEMNTSTVNSAREELMESRMRIESLSSQLSNLQKESRACLERIQELEDLLAKEKDNSR
RMLTDKEREMAEIRDQMQQQLNDYEQLLDVKLALDMEISAYRKLLEGEEERLKLSPSPSSRVTVSRASSSRSVRTTRGKRKRVD
VEESEASSSVSISHSASATGNVCIEEIDVDGKFIRLKNTSEQDQPMGGWEMIRKIGDTSVSYKYTSRYVLKAGQTVTIWAANAGVT
ASPPTDLIWKNQNSWGTGEDVKVILKNSQGEEVAQRSTVFKTTIPEEEEEEEEAAGVVVEEELFHQQGTPRASNRSCAIMGGDK
DCEMKRTTLDSPLGKLELSGCEQGLHRIIFLGKGTSAADAVEVPAPAAVLGGPEPLMQATAWLNAYFHQPEAIEEFPVPALHHP
VFQQESFTRQVLWKLLKVVKFGEVISYSHLAALAGNPAATAAVKTALSGNPVPILIPCHRVVQGDLDVGGYEGGLAVKEWLLAH
EGHRLGKPGLGSG[Hpep]*
LamB1 – Linker – SNAPf – Hpep

>pcDNA5/FRT/TO-TOMM20-SNAPf-Hpep

MVGRNSAIAAGVCGALFIGYCIYFDRKRRSDPNFKNRLRERRKKQKLAKERAGLSKLPDLKDAEAVQKFFLEEIQLGEELLAQGE
YEKGVDHLTNAIAVCGQPQQLLQVLQQTLPPPVFQMLLTKLPTISQRIVSAQSLAEDDVEGGSGDPPVGGDKDCEMKRTTLDSP
LGKLELSGCEQGLHRIIFLGKGTSAADAVEVPAPAAVLGGPEPLMQATAWLNAYFHQPEAIEEFPVPALHHPVFQQESFTRQVL
WKLLKVVKFGEVISYSHLAALAGNPAATAAVKTALSGNPVPILIPCHRVVQGDLDVGGYEGGLAVKEWLLAHEGHRLGKPGLGS
G[Hpep]*
TOMM20 – Linker – SNAPf – Hpep

>pcDNA5/FRT/TO-cpHaloΔ3-T2A-EGFP-P30-SNAPf-NLS3x (cpHaloΔ3 only)

M[cpHaloΔ3]GSGATNFSLLKQAGDVEENPGPMVSKGEELFTGVVPILVELDGDVNGHKFSVSGEGEG
DATYGKLTLKFICTTGKLPVPWPTLVTTLTYGVQCFSRYPDHMKQHDFFKSAMPEGYVQERTIFFKDDGNYKTRAEVKFEGDT
LVNRIELKGIDFKEDGNILGHKLEYNYNSHNVYIMADKQKNGIKVNFKIRHNIEDGSVQLADHYQQNTPIGDGPVLLPDNHYLST
QSALSKDPNEKRDHMVLLEFVTAAGITLGMDELYKSGRPPPPPPPPPPPPPPPPPPPPPPPPPPPPPPGGRSRSLEMDKDCEMK
RTTLDSPLGKLELSGCEQGLHRIIFLGKGTSAADAVEVPAPAAVLGGPEPLMQATAWLNAYFHQPEAIEEFPVPALHHPVFQQES
FTRQVLWKLLKVVKFGEVISYSHLAALAGNPAATAAVKTALSGNPVPILIPCHRVVQGDLDVGGYEGGLAVKEWLLAHEGHRLG
KPGLGGAPDPKKKRKVDPKKKRKVDPKKKRKEL
T2A – Linker – EGFP – SNAPf – NLS 3x

>pcDNA5/FRT/TO-cpHaloΔ3-T2A-gsg-Hpep11-gsg-EGFP-P30-SNAPf-NLS3x (N-terminal tagging)

M[cpHaloΔ3]GSGATNFSLLKQAGDVEENPGPGSGSKRDWREMFRLFRTGSGMVSKGEELFTGVVPILVELDGDVNGHKFSVSG
EGEGDATYGKLTLKFICTTGKLPVPWPTLVTTLTYGVQCFSRYPDHMKQHDFFKSAMPEGYVQERTIFFKDDGNYKTRAEVKF
EGDTLVNRIELKGIDFKEDGNILGHKLEYNYNSHNVYIMADKQKNGIKVNFKIRHNIEDGSVQLADHYQQNTPIGDGPVLLPDN
HYLSTQSALSKDPNEKRDHMVLLEFVTAAGITLGMDELYKSGRPPPPPPPPPPPPPPPPPPPPPPPPPPPPPPGGRSRSLEMDKD
CEMKRTTLDSPLGKLELSGCEQGLHRIIFLGKGTSAADAVEVPAPAAVLGGPEPLMQATAWLNAYFHQPEAIEEFPVPALHHPV
FQQESFTRQVLWKLLKVVKFGEVISYSHLAALAGNPAATAAVKTALSGNPVPILIPCHRVVQGDLDVGGYEGGLAVKEWLLAHE
GHRLGKPGLGGAPDPKKKRKVDPKKKRKVDPKKKRKEL*
T2A – Linker – EGFP – Hpep11 – SNAPf – NLS 3x

>pcDNA5/FRT/TO-cpHaloΔ3-T2A-EGFP-gsg-Hpep11-P30-SNAPf-NLS3x (C-terminal tagging)

M[cpHaloΔ3]GSGATNFSLLKQAGDVEENPGPMVSKGEELFTGVVPILVELDGDVNGHKFSVSGEGEGDATYGKLTLKFICTTGK
LPVPWPTLVTTLTYGVQCFSRYPDHMKQHDFFKSAMPEGYVQERTIFFKDDGNYKTRAEVKFEGDTLVNRIELKGIDFKEDGNI
LGHKLEYNYNSHNVYIMADKQKNGIKVNFKIRHNIEDGSVQLADHYQQNTPIGDGPVLLPDNHYLSTQSALSKDPNEKRDHMV
LLEFVTAAGITLGMDELYKGSGSKRDWREMFRLFRTSGRPPPPPPPPPPPPPPPPPPPPPPPPPPPPPPGGRSRSLEMDKDCEM
KRTTLDSPLGKLELSGCEQGLHRIIFLGKGTSAADAVEVPAPAAVLGGPEPLMQATAWLNAYFHQPEAIEEFPVPALHHPVFQQ
ESFTRQVLWKLLKVVKFGEVISYSHLAALAGNPAATAAVKTALSGNPVPILIPCHRVVQGDLDVGGYEGGLAVKEWLLAHEGHR
LGKPGLGGAPDPKKKRKVDPKKKRKVDPKKKRKEL*
T2A – Linker – EGFP – Hpep11 – SNAPf – NLS 3x

>pcDNA5/FRT/TO-cpHaloΔ3-T2A-EGFP-Hpep11[loopC_gsgx2]-P30-SNAPf-NLS3x (LoopC insertion)

M[cpHaloΔ3]GSGATNFSLLKQAGDVEENPGPMVSKGEELFTGVVPILVELDGDVNGHKFSVSGEGEGDATYGKLTLKFICTTGK
LPVPWPTLVTTLTYGVQCFSRYPDHMKQHDFFKSAMPEGYVQERTIFFKDDGNYKTRAEVKFEGDTLVNRIELKGIDFKEDGNI
LGHKLEYNYNSHNVYIMADKQKNGIKVNFKIRHNIEDGSGGSGSKRDWREMFRLFRTGSGGTGVQLADHYQQNTPIGDGPVLL
PDNHYLSTQSALSKDPNEKRDHMVLLEFVTAAGITLGMDELYKSGRPPPPPPPPPPPPPPPPPPPPPPPPPPPPPPGGRSRSLEM
DKDCEMKRTTLDSPLGKLELSGCEQGLHRIIFLGKGTSAADAVEVPAPAAVLGGPEPLMQATAWLNAYFHQPEAIEEFPVPALH
HPVFQQESFTRQVLWKLLKVVKFGEVISYSHLAALAGNPAATAAVKTALSGNPVPILIPCHRVVQGDLDVGGYEGGLAVKEWLL
AHEGHRLGKPGLGGAPDPKKKRKVDPKKKRKVDPKKKRKEL*
T2A – Linker – EGFP – Hpep11 – SNAPf – NLS 3x

>pcDNA5/FRT/TO-EGFP-DogTag[loopC_gsgx2]-P30-SNAPf-NLS3x (LoopC insertion)

MVSKGEELFTGVVPILVELDGDVNGHKFSVSGEGEGDATYGKLTLKFICTTGKLPVPWPTLVTTLTYGVQCFSRYPDHMKQHDF
FKSAMPEGYVQERTIFFKDDGNYKTRAEVKFEGDTLVNRIELKGIDFKEDGNILGHKLEYNYNSHNVYIMADKQKNGIKVNFKI
RHNIEDGTGGSGDIPATYEFTDGKHYITNEPIPPKGSGGTGVQLADHYQQNTPIGDGPVLLPDNHYLSTQSALSKDPNEKRDHM
VLLEFVTAAGITLGMDELYKSGRPPPPPPPPPPPPPPPPPPPPPPPPPPPPPPGGRSRSLEMDKDCEMKRTTLDSPLGKLELSGCE
QGLHRIIFLGKGTSAADAVEVPAPAAVLGGPEPLMQATAWLNAYFHQPEAIEEFPVPALHHPVFQQESFTRQVLWKLLKVVKFG
EVISYSHLAALAGNPAATAAVKTALSGNPVPILIPCHRVVQGDLDVGGYEGGLAVKEWLLAHEGHRLGKPGLGGAPDPKKKRKV
DPKKKRKVDPKKKRKEL*
Linker – EGFP – DogTag – SNAPf – NLS 3x
